# Development of a novel, pan-variant aerosol intervention for COVID-19

**DOI:** 10.1101/2021.09.14.459961

**Authors:** Robert H. Shoemaker, Reynold A. Panettieri, Steven K. Libutti, Howard S. Hochster, Norman R. Watts, Paul T. Wingfield, Philipp Starkl, Lisabeth Pimenov, Riem Gawish, Anastasiya Hladik, Sylvia Knapp, Daniel Boring, Jonathan M. White, Quentin Lawrence, Jeremy Boone, Jason D. Marshall, Rebecca L. Matthews, Brian D. Cholewa, Jeffrey W. Richig, Ben T. Chen, David L. McCormick, Romana Gugensberger, Sonja Höller, Josef M. Penninger, Gerald Wirnsberger

## Abstract

To develop a universal strategy to block SARS-CoV-2 cellular entry and infection represents a central aim for effective COVID-19 therapy. The growing impact of emerging variants of concern increases the urgency for development of effective interventions. Since ACE2 is the critical SARS-CoV-2 receptor and all tested variants bind to ACE2, some even at much increased affinity (see accompanying paper), we hypothesized that aerosol administration of clinical grade soluble human recombinant ACE2 (APN01) will neutralize SARS-CoV-2 in the airways, limit spread of infection in the lung and mitigate lung damage caused by deregulated signaling in the renin-angiotensin (RAS) and Kinin pathways. Here we show that intranasal administration of APN01 in a mouse model of SARS-CoV-2 infection dramatically reduced weight loss and prevented animal death. As a prerequisite to a clinical trial, we evaluated both virus binding activity and enzymatic activity for cleavage of Ang II following aerosolization. We report successful aerosolization for APN01, retaining viral binding as well as catalytic RAS activity. Dose range-finding and IND-enabling repeat-dose aerosol toxicology testing were conducted in dogs. Twice daily aerosol administration for two weeks at the maximum feasible concentration revealed no notable toxicities. Based on these results, a Phase I clinical trial in healthy volunteers can now be initiated, with subsequent Phase II testing in individuals with SARS-CoV-2 infection. This strategy could be used to develop a viable and rapidly actionable therapy to prevent and treat COVID-19, against all current and future SARS-CoV-2 variants.

**One Sentence Summary:** Preclinical development and evaluation of aerosolized soluble recombinant human ACE2 (APN01) administered as a COVID-19 intervention is reported.

## Introduction

Early in the COVID-19 pandemic, sequencing of SARS-CoV-2 enabled recognition of the high degree of homology with SARS-CoV and the identification of ACE2 as a candidate receptor for both viruses (*1,2*). A series of publications in early 2020 defined the molecular details regarding structural interactions between the receptor binding domain of SARS-CoV-2 and the ACE2 receptor (*3-6*). Blocking the Spike-ACE2 interaction provided a potential anti-SARS-CoV-2 therapeutic strategy and is the basis for virtually all successful vaccine designs (*7*). Proteins or peptides interacting with either of the binding partners have therapeutic potential and some very high affinity binders have been reported (*8*). Likewise, engineered antibodies can inhibit the Spike-ACE2 interaction, and several have received Emergency Use Authorization (EUA) as single agents or combinations from the FDA and EMA as systemic therapeutics (*9-11*). Recently, the EUA of a monoclonal antibody directed to the Spike protein was revoked for use as a single agent because of reduced activity against emerging viral variants (*12*). Use of recombinant ACE2 may prove to be a universal and robust therapeutic intervention, since all studied emerging SARS-CoV-2 variants continue to use ACE2 as the primary receptor. Importantly, although multiple entry receptors have been proposed, recent data have unequivocally shown that ACE2 is the essential SARS-CoV-2 receptor *in vivo* (*13*).

After the outbreak of the first SARS virus in 2003, soluble recombinant human ACE2 (APN01) was developed for systemic treatment of acute respiratory distress syndrome (ARDS) (*14,15*). In this indication, the catalytic activity of ACE2 in cleaving Ang II was exploited to reduce damage to the lung as observed in virus induced ARDS. Phase I and Phase II clinical trials demonstrated that APN01 had an acceptable safety profile and strongly reduced pathogenic Ang II levels (*16*). Our group first reported *in vitro* SARS-CoV-2 neutralizing activity of APN01 in cells and human organoids (*17*). Importantly, interactions between Spike proteins of multiple *variants of concern* and clinical grade soluble human recombinant ACE2 (APN01) have been demonstrated to be of considerably higher affinity. Moreover, APN01 can neutralize all tested SARS-CoV-2 variants of concern and variants of interest (see accompanying manuscript). The increased affinity to both APN01 and, by implication, endogenous ACE2 receptor is thought to contribute to the enhanced infectivity and transmissibility observed for several of these variants (*18*).

Besides the emerging escape mutants, one of the difficulties for widely applicable Spike blocking therapies for people exposed to the virus or at early stages of disease, especially against variants, is their intravenous application. APN01 has recently undergone a randomized Phase 2 clinical trial for treatment of severe COVID-19 using intravenous administration (<u>NCT04335136</u>, manuscript in preparation). We reasoned that direct introduction of APN01 into the airways could locally neutralize the virus, limiting the spread of infection, and thereby limit damage to the lung. To probe the potential for therapeutic activity following introduction into the airways, we tested intranasal treatment in a mouse model of SARS-CoV-2 infection (*13*) with clinical grade APN01. Results demonstrated strong protective activity, providing experimental proof of concept for direct APN01 administration into the airways. The critical path for such a novel intervention includes the development of an aerosol formulation of APN01 that retains both virus-binding activity and enzymatic activity. We report the successful development of inhalable APN01 and results of preclinical toxicology studies that support the safety of this intervention when administered by aerosol. These data pave the way for an inhalable universal early intervention strategy against all current and future SARS-CoV-2 variants.

## Results

### In vitro anti-SARS-CoV-2 activity of APN01

While anti-SARS-CoV-2 activity of soluble recombinant human ACE2 (APN01) has been reported previously (*17*), we first evaluated neutralizing activity of the cGMP produced, i.v. injectable APN01, in cell culture. As shown in Figure 1, one-hour exposure to concentrations of APN01 as low as 25 µg/ml completely neutralized SARS-CoV-2 as assessed in a four-day cytopathic effect (CPE) assay on Vero E6 cells. Thus, clinical grade ACE2 exhibits antiviral activity without any apparent toxic effects. See accompanying manuscript for affinity/avidity measurements and neutralization of variants of concern.

**Figure 1.**
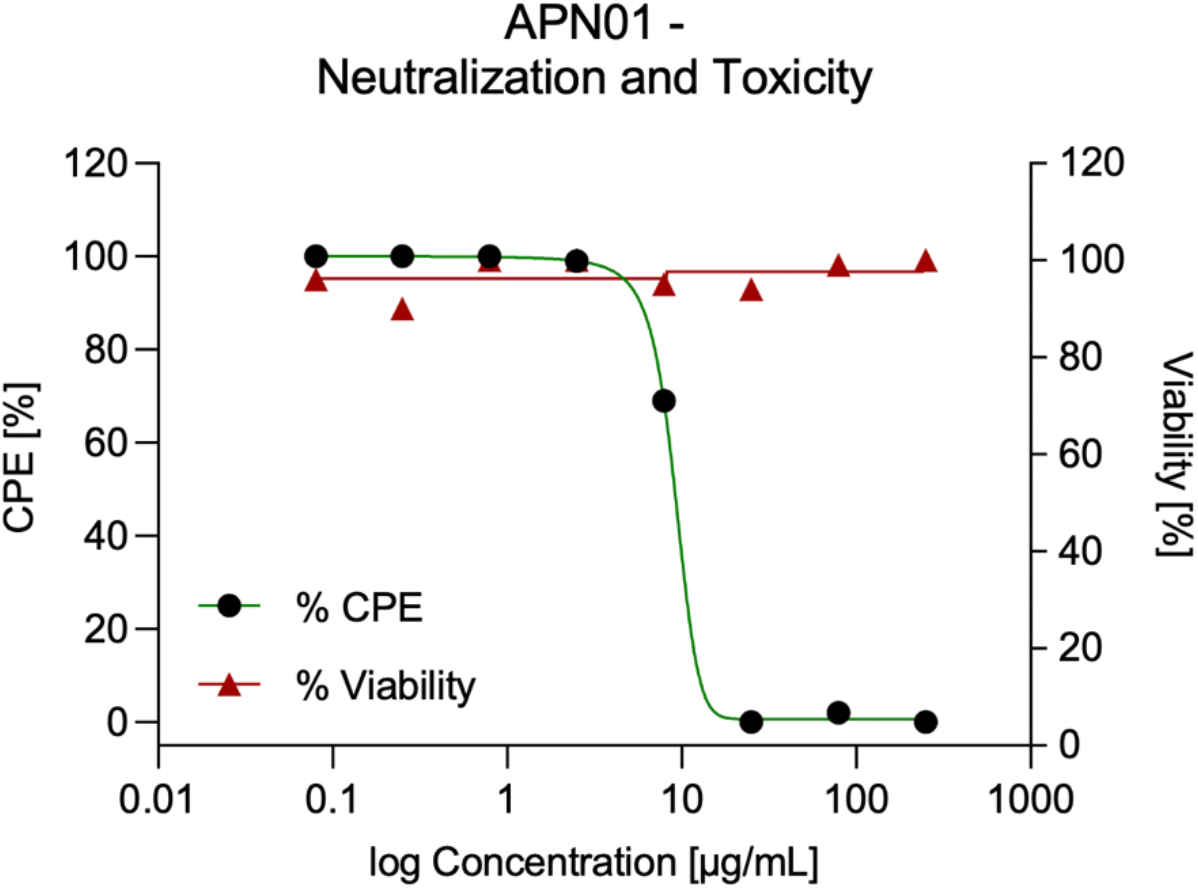
APN01 neutralization of SARS-CoV-2 in the absence of apparent cytotoxicity. Serial dilutions of APN01 were prepared in assay medium (MEM supplemented with 2% fetal bovine serum and 50 μg/mL gentamicin) and a suspension of SARS-CoV-2 (USA-WA1/2020) was added to assess neutralization. For assessment of APN01 on viability, assay medium without virus was added. After one-hour incubation at 37°C, the dilutions were transferred to wells containing Vero E6 target cells (Multiplicity Of Infection 0.001). Incubation was continued for four days and cell numbers were assessed with a neutral red endpoint. Cytopathic Effect (CPE) was calculated as the average optical density (OD) for replicate infected and treated wells divided by average control OD X 100 (expressed as a percentage). Viability was calculated as the average optical density (OD) for replicate uninfected and treated wells divided by average control OD X 100 (expressed as a percentage).

### APN01 protects from respiratory SARS-CoV-2 infections

We have recently developed a novel animal model that faithfully recapitulates SARS-CoV-2 infections and results in severe lung pathologies, weight loss, and, dependent on the mouse strain, in death of infected mice. This model is based on a mouse adapted SARS-CoV-2 virus (termed *maVie16*) and expression of ACE2 was found to be essential for infection and disease (*13*). We used this model to test whether application of clinical grade APN01 into the upper respiratory tract of SARS-CoV-2 *maVie16* infected mice might show clinical benefits (Figure 2A). Of note, SARS-CoV-2 *maVie16* can still effectively infect human cells, i.e. the *maVie16* Spike can bind to human ACE2. Whereas *maVie16* infection of Balb/c mice resulted in rapid weight loss, lung pathologies, and eventual death, intranasal APN01 delivery protected from lung damage and weight loss (Figure 2B). Treatment-related changes in body temperature did not reach statistical significance. Importantly, APN01 treatment resulted in 100% survival of the SARS-CoV-2 infected mice whereas all controls succumbed to the infection (infected mice surviving to day 5 were moribund when sacrificed). Increase in lung tissue weight, a correlate of severe infection, was also significantly reduced by intranasal APN01 treatment (Figure 2C). Thus, respiratory delivery of APN01 protects from SARS-CoV-2 infections.

**Figure 2.**
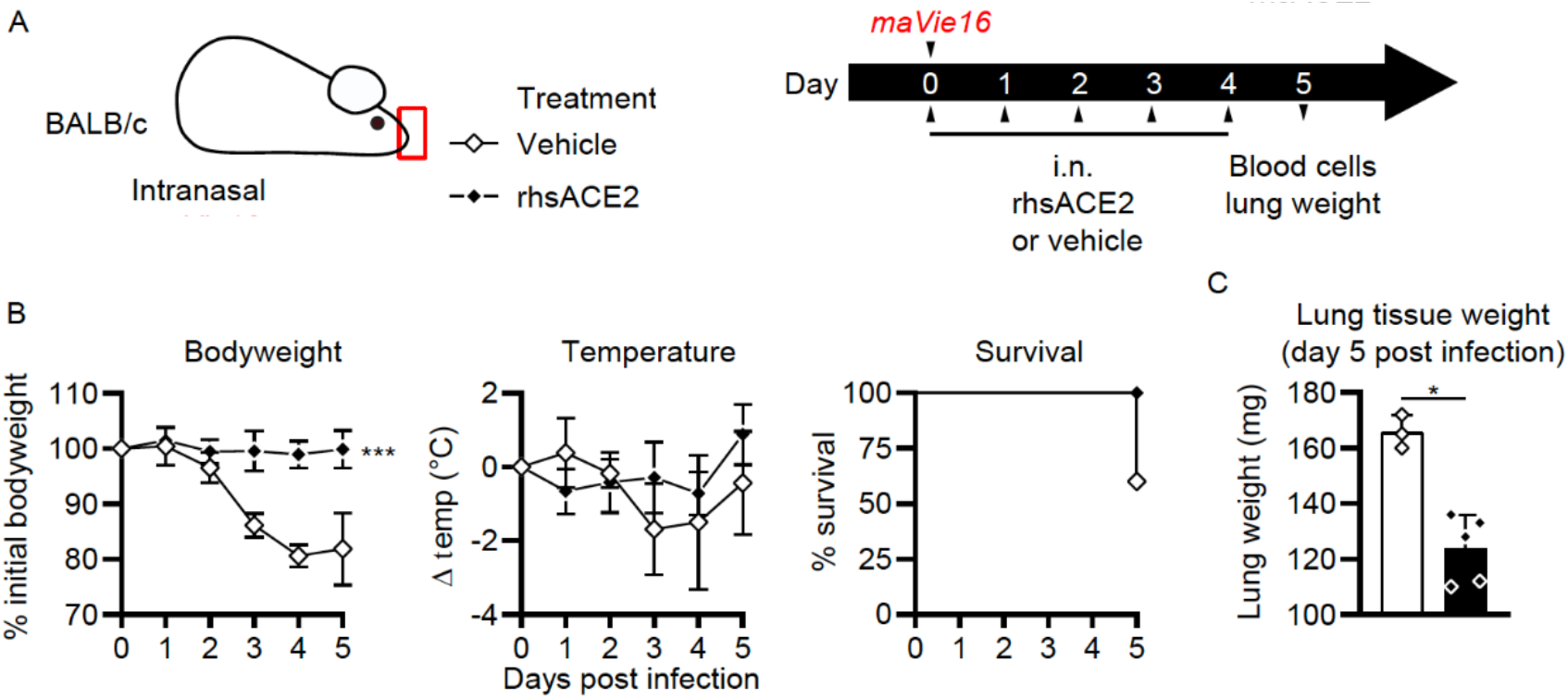
Intranasal APN01 protects from COVID-19 in a mouse adapted SARS-CoV-2 respiratory infection model. (A) Experimental outline for infection of BALB/c mice (n = 5 for both groups) with SARS-CoV-2 (strain *maVie16*) and daily intranasal treatment with APN01 for five days. (B) Body weight, temperature and survival curves for infected BALB/c mice treated with vehicle control or recombinant human soluble ACE2 (rhsACE2=APN01). Body weights and temperature were compared using mixed-effect analyses. Survival differences were analyzed using a Mantel-Cox test. (C) Lung tissue weight as assessed 5 days after infection of mice; data were analyzed with the Mann-Whitney test. Statistical significances are indicated by asterisks (p-value < 0.05: *; p-value < 0.001: ***).

### APN01 aerosolization

To aerosolize APN01 for preclinical studies in a way that can be scaled to future widespread and easy clinical use, we selected PARI LC PLUS nebulizers. These are widely available, effective and standardized devices already used in clinical applications. Aerosolized APN01 was collected using a custom fabricated condenser and analyzed for virus-binding activity and enzymatic activity for cleaving a fluorogenic substrate. Clinical grade APN01 is formulated for i.v. use at 5 mg/ml. Recognizing that *in vitro* anti-SARS-CoV-2 activity was observed at concentrations as low as 25 µg/ml for the reference USA-WA1/2020 virus (Figure 1) and even 10-20 times lower IC50/IC90 values for variants of concern and variants of interest (see accompanying paper), we aerosolized a range of concentrations ranging from 100 µg/ml to 2.5 mg/ml. The concentration of recovered APN01 was assessed by enzyme-linked immunosorbent assay (ELISA) measurements. Binding assays were conducted by coating plates with SARS-CoV-2 Spike RBD-Fc fusion protein and assessing APN01 binding (Figure 3). Importantly, APN01 binding to SARS-CoV-2 Spike protein was almost identical after nebulization. Similar results were obtained in replicate assays for APN01 nebulized at 2.5 mg/ml and 0.1 mg/ml (Supplemental Table 1).

**Figure 3.**
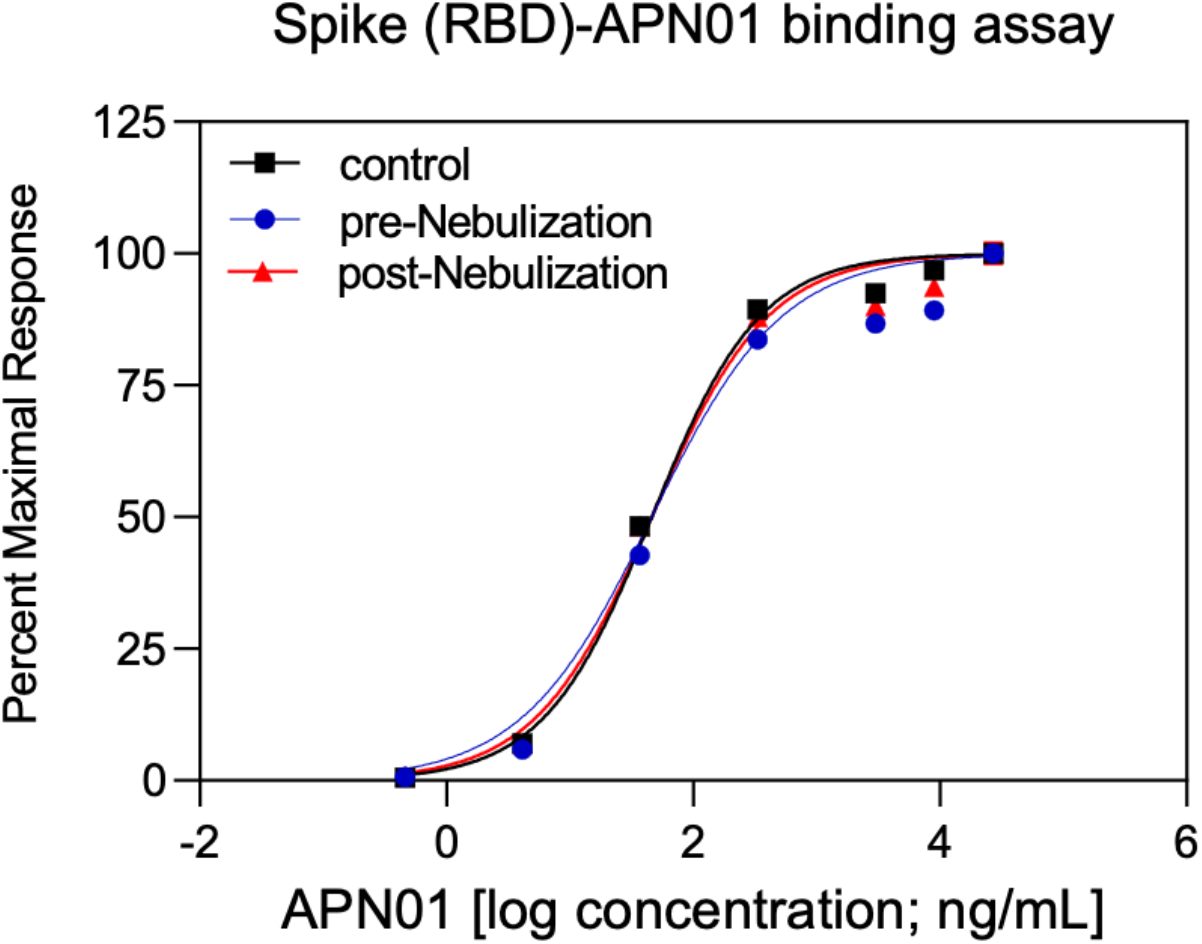
APN01 binding to the SARS-Cov-2 RBD is not altered by nebulization. Binding of increasing doses of APN01 to plate-immobilized RBD domain was assessed by ELISA. Curves depict pre-nebulization samples of APN01 (material remaining as liquid in the nebulizer cup), APN01 collected after nebulization (post-nebulization), and infusable non-treated control APN01. For this experiment, APN01 was aerosolized at 2.5 mg/ml.

To test for a potential effect of nebulization on the enzymatic activity of APN01, cleavage of a fluorogenic peptide substrate was assessed. Kinetic analysis was performed at multiple concentrations of APN01 and expressed as the change in Relative Fluorescence Units per minute per ng [(ΔRFU/min)/ng]. Results for a representative assay are illustrated in Figure 4. Enzymatic activity for cleaving the fluorescently labeled peptide substate by APN01 aerosolized at lower concentration was also not affected. Supplemental Table 2 presents results from replicate experiments with APN01 nebulized at 2.5 and 0.1 mg/ml. In summary, aerosolization of APN01 did not affect the structural integrity of APN01 in terms of its ability to bind SARS-CoV-2 Spike RBD nor its enzymatic activity.

**Figure 4.**
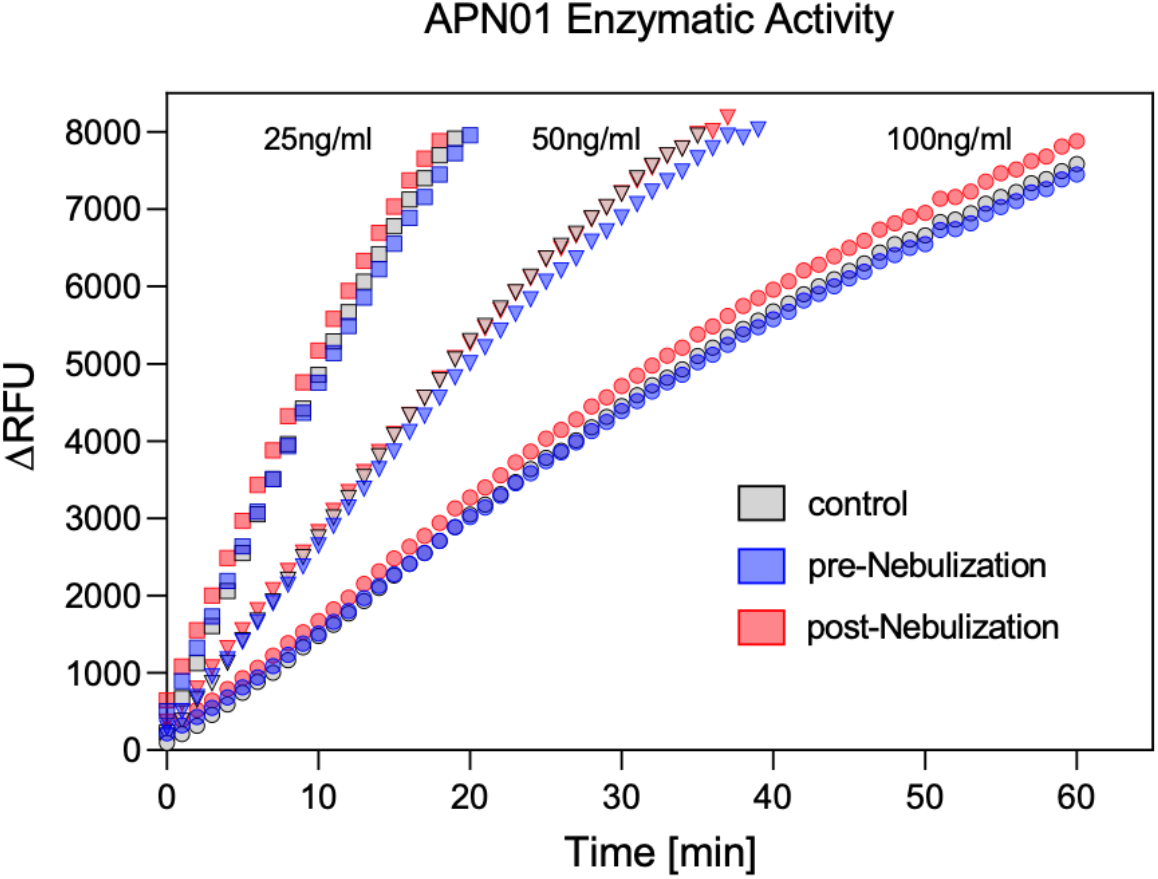
APN01 enzymatic activity is not altered by nebulization. Three samples were tested as defined in Figure 3 legend. Samples were diluted to APN01 concentrations of 25 ng/ml, 50 ng/ml, and 100 ng/ml and assayed for enzymatic function by cleavage of quenched fluorogenic MAPL-DNP substrate and subsequent measurement of fluorescence activity. Data are plotted as ΔRFU (change in relative fluorescence units) vs. time (min). One representative experiment out of 4 biological replicates is shown.

### Toxicologic assessment of assessment of aerosolized APN01

We next performed a comprehensive evaluation of the possible toxicity of twice daily inhalation administration of APN01 aerosols to beagle dogs for 14 consecutive days. Goals of this study, which was designed to support regulatory filings, included: (1) characterization of toxicities associated with repeat-dose inhalation exposure to APN01 aerosols; (2) identification of sensitive target tissues of inhaled APN01; (3) characterization of serum levels and toxicokinetics (TK) of inhaled APN01; and (4) identification of a No Observed Adverse Effect Level [NO(A)EL]. The study design and different test cohorts are summarized in Table 1.

**Table 1.** Toxicology design. Nebulized APN01 was administrated twice a day for 14 consecutive days at a low APN01 concentration (0.019 mg/L), a mid concentration (0.038 mg/L), and a high APN01 concentration (0.075 mg/L) for each single administration. The high APN01 dose was demonstrated to be the maximum feasible concentration (MFC), obtained by aerosolizing the i.v. formulation in a dose range-finding study.

| Group | Number of Dogs (M + F) | Agent | Number and Duration of Daily Exposures | Number of Exposure Days | Target APN01 Concentration in Test Atmosphere (mg/L) |
| --- | --- | --- | --- | --- | --- |
| 1 | 3 + 3 | Saline (Control) | 2 x 60 minutes | 14 | 0 |
| 2 | 3 + 3 | Vehicle (Control) | 2 x 60 minutes | 14 | 0 |
| 3 | 3 + 3 | APN01 – Low | 2 x 60 minutes | 14 | 0.019 |
| 4 | 3 + 3 | APN01 – Mid | 2 x 60 minutes | 14 | 0.038 |
| 5 | 3 + 3 | APN01 - High | 2 x 60 minutes | 14 | 0.075 |

For treatment of experimental animals, multiple PARI LC PLUS nebulizers were multiplexed through a distribution plenum with hoses connected to oronasal masks. Analytical data (obtained by HPLC analysis of filters) demonstrated that test aerosols consistently achieved target exposure concentrations. Characteristics of the generated APN01 aerosols are summarized in Table 2. Particle size distribution data showed that the generated particles were within the respirable size range for dogs (*19*) and met targets for both median mass aerodynamic diameter (MMAD) and geometric standard deviation (GSD).

**Table 2.** Characterization of APN01 aerosols. Samples were collected at morning (AM) and afternoon (PM) exposures and characterized for APN01 concentrations to enable dose calculation and assess particle sizes to confirm respirability. GSD, geometric standard deviation; MMAD, median mass aerodynamic diameter.

| Group<br>(Treatment) | Sample<br>Collection<br>Time Point | Analytical<br>Concentration<br>(mg/L) <sup>a,b,c</sup> | Particle Size Distribution |  |
| --- | --- | --- | --- | --- |
|  |  |  | MMAD <sup>a,d</sup><br>(µm) | GSD<br>Range<sup>d</sup> |
| 1<br>(Saline Control) | AM | 0.000 ± 0.0000 | 1.06 ± 0.113 | 2.25-2.36 |
|  | PM | 0.000 ± 0.0000 |  |  |
| 2<br>(Vehicle Control) | AM | 0.000 ± 0.0000 | 1.36 ± 0.078 | 2.20-2.56 |
|  | PM | 0.000 ± 0.0000 |  |  |
| 3<br>(APN01; 0.019 mg/L) | AM | 0.017 ± 0.0014 | 1.57 ± 0.566 | 1.53-2.28 |
|  | PM | 0.017 ± 0.0028 |  |  |
| 4<br>(APN01; 0.038 mg/L) | AM | 0.034 ± 0.0034 | 1.92 ± 0.205 | 1.80-2.41 |
|  | PM | 0.035 ± 0.0024 |  |  |
| 5<br>(APN01; 0.075 mg/L) | AM | 0.078 ± 0.0062 | 2.00 ± 0.290 | 1.87-1.95 |
|  | PM | 0.082 ± 0.0058 |  |  |
| <sup>a</sup> Mean ± standard deviation |  |  |  |  |
| <sup>b</sup> N = 15 (each study group was exposed for 14 days; however, due to staggered dosing for the two sexes, aerosol samples were collected over 15 days) |  |  |  |  |
| <sup>c</sup> Analytical concentration calculated based on HPLC analysis results:<br>mg/filter HPLC result/sample volume (L) |  |  |  |  |
| <sup>d</sup> N = 2 (one measurement per study week) |  |  |  |  |

The inhaled dose of APN01 following one hour exposure was calculated from the aerosol concentration, average volume inhaled (Minute Volume in L/min), exposure time and animal body weights. Calculated inhaled APN01 levels for each group after a single exposure (Study Day 1) and after repeat-dose exposure (after the first exposure on Study Day 11) are shown in Table 3. Inhaled doses were concentration dependent and consistent over the treatment interval. Even dosed at the lowest concentration, the inhaled dose was >0.3 mg/kg/exposure, far exceeding the minimum virus neutralizing activity of APN01.

**Table 3.** Calculated inhaled dose levels. At all concentrations, the inhaled APN01 dose exceeded SARS-CoV-2 neutralizing concentrations. This was the case for both the initial administration and after repeated exposures.

| Group (Treatment) | Sex | Test Atmosphere APN01 Concentration <sup>a</sup> (mg/L) |  | Minute Volume (L/min) |  | Mean Body Weight (kg) |  | Calculated Inhaled APN01 Dose (mg/kg/Exposure) <sup>b</sup> |  |
| --- | --- | --- | --- | --- | --- | --- | --- | --- | --- |
|  |  | Post Single Exposure<sup>c</sup> | Post Repeated Exposures<sup>d</sup> | Post Single Exposure<sup>c</sup> | Post Repeated Exposures<sup>d</sup> | Post Single Exposure<sup>c</sup> | Post Repeated Exposures<sup>d</sup> | Post Single Exposure<sup>c</sup> | Post Repeated Exposures<sup>d</sup> |
| 1 (Saline Control) | M | 0 | 0 | 3.0153 | 2.9915 | 9.24 | 9.15 | 0.0 | 0.0 |
|  | F | 0 | 0 | 2.5442 | 2.5470 | 7.49 | 7.50 | 0.0 | 0.0 |
| 2 (Vehicle Control) | M | 0 | 0 | 2.8906 | 2.9650 | 8.77 | 9.05 | 0.0 | 0.0 |
|  | F | 0 | 0 | 2.4337 | 2.4864 | 7.09 | 7.28 | 0.0 | 0.0 |
| 3 (APN01; 0.019 mg/L) | M | 0.019 | 0.017 | 2.8773 | 2.9332 | 8.72 | 8.93 | 0.38 | 0.34 |
|  | F | 0.016 | 0.013 | 2.3585 | 2.3529 | 6.82 | 6.80 | 0.33 | 0.27 |
| 4 (APN01; 0.038 mg/L) | M | 0.027 | 0.035 | 2.9199 | 2.8986 | 8.88 | 8.80 | 0.53 | 0.69 |
|  | F | 0.026 | 0.035 | 2.3753 | 2.3725 | 6.88 | 6.87 | 0.54 | 0.73 |
| 5 (APN01; 0.075 mg/L) | M | 0.068 | 0.078 | 2.9066 | 2.9119 | 8.83 | 8.85 | 1.34 | 1.54 |
|  | F | 0.073 | 0.080 | 2.4947 | 2.5470 | 7.31 | 7.50 | 1.49 | 1.63 |

<sup>a</sup> Determined by HPLC analysis
<sup>b</sup> Inhaled dose calculated as: **Dose (mg/kg/day) = (C × MV × T) / BW**, where C is the test atmosphere concentration (mg/L) per group (first of two daily exposures); MV is the group mean minute volume for the appropriate sex; calculated as (L/min) = 0.499 × BW<sup>0.809</sup> \*; T is the duration of exposure (60 minutes); BW is the group mean body weight for the appropriate sex. \* Bide, RW, et al. *J. Appl. Toxicol.* **20**, 273–290 (2000)
<sup>c</sup> Study Day 1 after the first of two daily exposures.
<sup>d</sup> Study Day 11 after the first of two daily exposures (21 total exposures); data from this time point were used since this was the last available non-fasted body weight (twice weekly weighing schedule).

We also tested whether inhalation of APN01 might result in dispersion of APN01 outside the respiratory system. Systemic exposure [defined as serum levels of APN01 above the limit of quantitation (LOQ)] was very low in dogs in the low dose and mid dose groups as assessed on Days 1 and 14; in both groups, serum levels of APN01 were below the LOQ (0.5 ng/mL) in most animals at most time points (Table 4). In the high dose group, serum levels of APN01 on Days 1 and 14 were above the LOQ in 5 out of 6 dogs at all time points, tested after 0.5 hr following inhalation. Although interanimal variability was substantial, mean C_max_ (pooled across both sexes) in the high dose group was approximately 8 ng/mL on both Day 1 and Day 14. The systemic exposure to APN01 following aerosol administration was far below the levels attained following i.v. administration in the clinic where plasma concentrations have been reported in excess of 1000 ng/mL (*20*).

**Table 4.** Serum levels of inhaled APN01 and toxicokinetic parameters. Dog serum samples were analyzed for levels of APN01 using ELISA. At scheduled blood sampling times (Days 1 and 14), serum was sampled from individual animals in Groups 3-5 and subsequently used to model toxicokinetic (TK) parameters. Reported TK parameters are: time at which maximum concentration was observed (T_max_), maximum concentration observed (C_max_), and area under the concentration curve at six hours (AUC_6hr_). Terminal phase parameters [e.g., T_1/2_] could not be calculated reliably for any of the profiles due to a lack of amenable data in the terminal phase of the profiles. Data for males and females are shown.

| Group | Target Test<br>Atmosphere APN01<br>Concentration (mg/L) | Estimated Mean Inhaled<br>APN01 Dose<br>(mg/kg) | Mean / *Median* Group TK Parameter |  |  |  |
| --- | --- | --- | --- | --- | --- | --- |
|  |  |  | n | T<sub>max</sub><br>(hr) | C<sub>max</sub><br>(ng/mL) | AUC<sub>6hr</sub><br>(hr*ng/mL) |
| MALES |  |  |  |  |  |  |
| Day 1 |  |  |  |  |  |  |
| 3 | 0.19 | 0.38 | 2 | 1 | 1.26 | -- |
| 4 | 0.038 | 0.53 | 3 | 2 | 1.92 | 16.3 |
| 5 | 0.075 | 1.34 | 3 | 1 | 4.38 | 21.1 |
| Day 14 |  |  |  |  |  |  |
| 3 | 0.19 | 0.34 | 1 | 1 | 1.04 | -- |
| 4 | 0.038 | 0.69 | 3 | 0.5 | 1.44 | -- |
| 5 | 0.075 | 1.54 | 3 | 1 | 7.96 | 38.0 |
| FEMALES |  |  |  |  |  |  |
| Day 1 |  |  |  |  |  |  |
| 3 | 0.19 | 0.33 | 3 | 6 | 3.63 | 20.1 |
| 4 | 0.038 | 0.54 | 2 | 1 | 1.64 | -- |
| 5 | 0.075 | 1.49 | 3 | 6 | 11.5 | 54.0 |
| Day 14 |  |  |  |  |  |  |
| 3 | 0.19 | 0.27 | 2 | 0.25 | 3.73 | 13.5 |
| 4 | 0.038 | 0.73 | 2 | 0.38 | 2.26 | 9.87 |
| 5 | 0.075 | 1.63 | 3 | 0.5 | 7.50 | 28.9 |

Toxicology endpoints also included mortality/moribundity observations; clinical observations for signs of toxicity; physical examinations; heart rate and blood pressure measurements; body weight measurements; food consumption measurements; ophthalmic examinations; electrocardiographic evaluations; respiratory function evaluations; measurements of blood oxygen saturation and pH; neurotoxicity evaluations (functional observational battery [FOB]); clinical pathology assessments (clinical chemistry, hematology, coagulation, and urinalysis); quantitation of serum drug levels; limited modeling of serum toxicokinetics (TK); gross pathology at necropsy; organ weights; and microscopic evaluation of tissues. See Supplemental Table 3 for a listing of toxicology parameters studied.

No early deaths occurred during the study and no gross clinical signs of toxicity were seen in any study animal. Inhalation administration of APN01 aerosols had no effects on body weight, food consumption, clinical pathology parameters, heart rate, blood pressure, electrocardiography, blood oxygen saturation, blood pH, FOB parameters, or ophthalmology. Respiratory function evaluations (respiratory rate, tidal volume and minute volume) were inconclusive due to excitement and/or panting exhibited by study animals during measurement periods. Organ weights were comparable in all study groups. No gross or microscopic pathology was linked to APN01 administration. Moreover, no evidence of systemic or organ-specific toxicity was identified in any dog receiving twice daily 60-minute exposures to APN01 aerosols at all target concentrations of 0.019, 0.038, or 0.075 mg/L for fourteen consecutive days. On this basis, the NO(A)EL was determined to be the highest dose investigated, i.e. 0.075 mg/L. Thus, in agreement with toxicologic assessments of systemic exposure of APN01 in rodent and non-rodent species, APN01 exhibited an excellent tolerability and safety profile upon administration as an aerosol.

## Discussion

Animal modeling conducted in this study with a mouse adapted SARS-CoV-2 strain has provided critical preclinical evidence for the efficacy of APN01 to prevent COVID-19 symptoms when delivered directly to the sites of respiratory infection via intranasal administration. These data are in agreement with the efficacy of ACE2 derived peptides delivered intranasally in a hamster model of COVID-19 (*8*), as well as novel lipopeptides, designed to inhibit viral entry, delivered as a nasal spray in a ferret model (*21*), all of these data pointing the way for early intervention using airway administered therapeutics. Approaches for limiting viral entry early in the course of infection via ACE2 based drugs may be complemented by approaches delivering antiviral drugs such as remdesivir and interferon β directly into the airways and lungs. Indeed combinations of remdesivir and interferon β are planned for clinical studies in the near future (<u>NCT04647695</u>).

As a COVID-19 intervention, APN01 offers key advantages, especially for treatment of emerging variants and APN01 exhibits markedly increased affinities/avidities to the variants RBD/Spike and enhanced neutralization activity (please see accompanying manuscript). Mutations detected in variants of concern and variants of interest cluster in the viral Spike protein and particularly in the receptor binding domain. Although these variants can in part escape neutralization by vaccines, convalescent plasma, and multiple therapeutic antibodies, all of these variants continue to use ACE2 as primary receptor. Importantly, our recent data for the first time also show that ACE2 deficient mice no longer exhibit signs of COVID-19 in our mouse adapted SARS-CoV-2 *maVie16* infection model (18), demonstrating that ACE2 is essential for SARS as well as SARS-CoV-2 infections. The mouse adapted SARS-CoV-2 *maVie16* strain bears two key mutations in the RBD to allow for mouse ACE2 receptor binding but remains susceptible to APN01. Thus, our novel and severe mouse COVID-19 model allowed us to assess the therapeutic activity of APN01 directly administered to the respiratory tree. Indeed, treatment with APN01 at the time of infection prevented any COVID-19 pathologies (lung), clinical symptoms (weight loss) and, importantly, completely protected from death. Similarly, intranasal therapy with murine ACE2, produced in exactly the same manner as APN01, completely prevented the SARS-CoV-2 *maVie16* infection and all mice remained healthy (*13*), consistent with the APN01 studies. Since in contrast to therapeutic MoAbs or natural Abs (22,23) no virus escape mutants should occur in binding to ACE2, APN01 could be used as an inhalable universal therapy against all current and future SARS-CoV-2 variants. The results presented here support the high therapeutic potential and feasibility of the administration of APN01 into the airways and lungs of patients for treatment of SARS-CoV-2 infection. Aerosolized APN01 retains virus binding and enzymatic activities. Furthermore, the aerosol generated using a commercial and widely used nebulizer has a particle size distribution consistent with delivery throughout the respiratory tract and could be delivered repeatedly at high doses to dogs without notable toxicities. Thus, APN01 aerosol administration should deliver effective antiviral therapy to the airways.

A limitation of our toxicology results is that only a single species of experimental animals is included. Additional confidence in the results could be generated by including non-human primates in the analysis. Of note, mice also did not show any signs of pathologies when they received APN01 or mouse soluble ACE2 into their respiratory system for 5 day efficacy studies. Moreover, the prior experience in clinical administration of APN01 in severe COVID-19 patients via the i.v. route, without serious adverse events (manuscript in preparation), supports moving ahead with Phase I testing using a conservative dose escalation strategy. A starting clinical dose of ¼ the maximum feasible concentration for 15 minutes provides an estimated 100-fold safety margin assuming 100% aerosol deposition when compared to the NO(A)EL observed in dogs. Escalation to twice per day and then increasing concentrations of ½ to the maximum feasible concentration is the anticipated design. A Phase I trial for safety and tolerability of aerosolized APN01 in healthy volunteers is currently underway (NCT number pending) which will be followed by Phase II trials in individuals infected with SARS-CoV-2. The latter trials will use viral clearance as the primary endpoint with severe disease and hospitalization as secondary endpoints.

In summary, our study demonstrates both the potent preclinical activity of locally administered APN01 in a mouse model of COVID-19 as well as the feasibility and excellent safety profile of APN01 delivered as an aerosol in a toxicologic assessment in dogs. These data, along with the data provided in our companion manuscript showing activity against numerous emerging variants, highlight the potential of aerosolized APN01 to serve as a potent pan-SARS-CoV-2 therapeutic.

## Materials and Methods

### *In vitro* anti-SARS-CoV-2 neutralizing activity of APN01

Serial dilutions of APN01 were prepared in assay medium (MEM supplemented with 2% fetal bovine serum and 50 μg/ml gentamicin) and a suspension of SARS-CoV-2, USA-WA1/2020 was added to assess neutralization. For assessment of APN01 on viability, assay medium without virus was added. After one-hour incubation at 37°C, the dilutions were transferred to wells containing Vero E6 target cells (MOI 0.001). Incubation was continued for four days and cell numbers were assessed with a neutral red endpoint. Testing was performed under contract at the Institute for Antiviral Research, Utah State University, Logan, UT. All of the experiments reported here used clinical grade recombinant soluble human ACE2 (APN01).

### Activity of APN01 in a mouse model of SARS-CoV-2 infection

Ten-week-old BALB/c mice (Charles River) received daily intranasal treatments with 100 µg APN01 or the respective dilution of vehicle in endotoxin-free PBS (Gibco). The first dose was given as a mix with 1 × 10^5^ TCID_50_ of *maVie16*, a mouse-adapted SARS-CoV-2 virus (18), in 50 µl, the remaining doses were administered in 40 µl endotoxin-free PBS to isoflurane-anesthetized animals. All experiments involving SARS-CoV-2 or its derivatives were performed in Biosafety Level 3 (BSL-3) facilities at the Medical University of Vienna and performed after approval by the institutional review board of the Austrian Ministry of Sciences (BMBWF-2020-0.253.770) and in accordance with the directives of the EU.

### Recovery of aerosolized APN01

An adaptor was connected to the mouthpiece port of the PARI LC PLUS nebulizer, secured to a ring stand, to route the aerosol into a custom fabricated condenser consisting of a 4L polypropylene graduated cylinder lined with downward-spiraling C-Flex tubing (Cole Palmer). The cylinder was filled with an ice water bath and the exit tubing was then fed through a hole into a 50 mL conical collection tube. The nebulizer was connected to a PARI Vios PRO Compressor according to the manufacturer’s instructions. For Trials 1-2, APN01 (lot ACE20620-B) was diluted to a volume of 4 ml at 2.5 mg/ml, transferred to the nebulizer cup, and nebulized for 24 min. For Trials 3-4, APN01 was diluted to a volume of 4 ml at 0.1 mg/ml and nebulized for 24 min. This resulted in a range of recovery from all trials of 0.570-1.420 ml APN01 volume in the collection tube, corresponding to 15.4-38.4% of the starting volume. Three samples were tested: diluted APN01 sample before addition to the nebulizer cup (Control), Unnebulized sample remainder in the nebulizer cup (Pre), and sample obtained from the collection tube after condensation (Post).

### ELISA to assess binding of APN01 to the SARS-CoV-2 receptor binding domain (RBD)

96-well ELISA plates (Thermo-Fisher Scientific) were coated with 2 µg/ml SARS-CoV-2 Spike RBD-Fc protein (Sino Biological) in coating buffer (0.1 M carbonate-bicarbonate pH 9.4). Plates were sealed with adhesive plate seals and incubated overnight at 4 °C, then washed 3X with wash buffer (1X PBS + 0.05% Tween 20). Plates were blocked with blocking buffer (1X PBS + 1% BSA), sealed, and incubated for 1 h at r.t. APN01 samples recovered from the nebulization runs were diluted in blocking buffer to 27 µg/ml, as quantified with hACE2 ELISA measurements. Further dilutions were conducted using blocking buffer for a seven-point standard curve. All 7 standards and a blank were added to the ELISA plate in triplicate. Plates were covered and incubated for 1 h at r.t., then washed 5X with wash buffer. Biotinylated goat anti-human ACE2 (R&D Systems) diluted in blocking buffer was added to plates at 0.4 µg/ml followed by a 1 h incubation at r.t. Plates were washed 5X with wash buffer. Streptavidin-HRP (Thermo-Fisher Scientific) diluted in blocking buffer was added to plates at 0.1 µg/mL followed by a 30 min incubation at r.t., then washed 3X with wash buffer. TMB substrate solution (SeraCare) was added followed by a 6 min incubation in the dark at r.t. Finally, stop solution 1 M H_2_SO_4_ (J.T. Baker, Inc.) was added and plates were gently shaken for 5 s to mix and then absorbance (O.D.) was measured at 450/620 nm on a SpectraMax M5 (Molecular Devices) instrument and data were processed by SoftMax Pro 6.4 software (Molecular Devices). Blank well readings were subtracted from sample dilution readings, and nonlinear regression to derive EC_50_ values was performed through a 4-parameter logistic equation by GraphPad Prism v.8 plotting OD450/620 versus APN01 (ng/ml).

### Angiotenin II cleavage assay

Substrate was prepared by reconstituting MAPL-DNP (Anaspec) in DMSO to make 1 mM stock with gentle mixing by inversion to ensure dissolution. This solution was diluted in assay buffer (10 µM ZnCl_2_, 50 mM MES, 300 mM NaCl, 0.01% Brij L23, pH 6.5) to 1 mM and then further diluted 1:5 in assay buffer to prepare a 0.2 mM working solution of MAPL-DNP. The substrate was prewarmed in an oven at 37 °C prior to addition to the sample. APN01 samples recovered from the nebulization runs were diluted in assay buffer to 1.0 µg/ml, as quantified with hACE2 ELISA measurements. Further dilution in assay buffer was conducted to yield 100 ng/ml, 50 ng/ml, and 25 ng/ml solutions at volumes of 1 ml each. The three APN01 dilutions of each sample were loaded into the assay plate (Greiner) in quadruplicate in the center of the plate, avoiding the outer edges, while assay buffer blanks were loaded in columns 2 and 11. Prewarmed 0.2 mM MAPL-DNP substrate was added to all wells in the plate starting with row A and contents in all rows were mixed by pipetting. Plates were read immediately for fluorescence (320/420 nm) in kinetic mode on a SpectraMax M5 (Molecular Devices) instrument at 37 °C for 1 h taking readings at 1 min intervals. Background-subtracted and averaged sample values (RFU) were plotted in Excel versus time, and slopes were determined (ΔRFU/min) for each sample. Data were converted to (ΔRFU/min)/ng APN01, where ΔRFU/min is the slope and ng APN01 was determined by multiplying the ng/mL concentration by 0.05 for the 50 µl of sample used per well.

#### ELISA to assess APN01 in dog plasma

Serum samples were analyzed for levels of APN01 using a commercial kit (Human ACE2 ELISA Kit PicoKine™; SKU EK0997; Boster Bio, Pleasanton, CA).

### HPLC quantitation of APN01 on filters

The aerosol mass concentration in each oronasal inhalation exposure system was determined by collecting the aerosol on glass-fiber filters. Samples were collected at a constant flow rate equal to the port flow of the delivery tube, and the total volume of air samples was measured by a dry-gas meter. One aerosol sample per dose level was collected during each exposure. Filter samples were extracted with phosphate buffered saline (PBS) and stored refrigerated (approximately 4°C). Filter samples were analyzed for levels of APN01 using high performance liquid chromatography (HPLC) with UV wavelength detection according to a validated method. During method validation, quality control (QC) samples prepared in PBS at target APN01 concentrations of 49 and 245 μg/mL were demonstrated to be stable for at least 22 days when stored refrigerated (111% and 104% of the original value, respectively) and for at least 2 days when stored at room temperature (111% and 103% of the original value, respectively). For instrument calibration, primary standard solutions of APN01 with target concentrations of 490 μg/mL were prepared by dilution of 0.5 mL of the test article formulation (4.9 mg/mL ACE2 content per the Certificate of Analysis) in PBS in a 5 mL volumetric flask and filling to volume. Standard curve calibrators were prepared at target concentrations of 29.4, 44.1, 58.8, 117.6, 176.4, 264.6 and 352.8 μg/mL by diluting the 490 μg/mL primary standards with PBS. QC samples of APN01 in PBS were prepared at low (50 μg/mL) and high (250 μg/mL) target concentrations and were analyzed along with the filter samples. Calibrator, QC and processed filter samples were analyzed by HPLC using the following equipment and conditions: HPLC System: Waters Alliance 2695; HPLC Detector: Waters 2487 Dual λ Absorbance Detector; Data System: Waters Empower3; HPLC Column: Agilent PLRP-S (reversed phase); 300 Å; 250 × 4.6 mm; 8 μm; Column Temperature: 25° C; Sample Temperature: 4° C; Injection Volume: 5 μL; Flow Rate: 1.0 mL/minute; Mobile Phase (MP): A: 0.1% trifluoroacetic acid in ASTM type I water B: 0.1% trifluoroacetic acid in acetonitrile. The retention time of APN01 was approximately 5 to 6 minutes. The calibration curve was calculated from the linear regression of the calibrators’ peak areas versus their respective concentrations. The concentrations of test article in the processed filter samples were determined from each sample’s peak area using the linear regression parameters derived from the calibration curves and correcting the resulting concentration by multiplying by the appropriate dilution factor, as applicable. A representative chromatogram is shown in Supplemental Figure 1.

### Aerosol Particle Size Distribution

Aerosol particle size distribution was determined twice per group during the study by collecting size-segregated aerosol samples using a 10-stage quartz crystal microbalance (QCM) cascade impactor (California Measurements Inc.; Sierra Madre, CA). The aerosol output from one port of the exposure system was connected to the QCM and was sampled at least once per inhalation exposure level. The mass median aerodynamic diameter (MMAD) and geometric standard deviation (GSD) of the test aerosol were calculated from the mass accumulated on each collection stage of the QCM by using a validated computer program (QCMSIZE) that was developed at IITRI.

### Toxicology Studies

Toxicology studies were conducted under Good Laboratory Practices. Animal studies were performed in full compliance with the Animal Welfare Act and in accordance to the NIH Guide for the Care and Use of Laboratory Animals.

## Acknowledgments

We thank Beverly Smolich of CCS Associates, San Jose, CA for advice on regulatory requirements for the aerosol intervention. We want to also thank all members of our laboratory and clinical groups for critical input.

## Funding

This work was supported by the National Cancer Institute through contracts with Leidos Biomedical Research (Contract Number 75N91019D00024 Task Order 75N91020F00003), MRIGlobal (Contract Number 75N91018D00026 Task Order 75N91019F00129), and IITRI (Contract Number 75N91019D00013 Task Order 75N91020F00002) and by NIH Grant UL1-TR-003017-03 to R.A.P. Funding for J.M.P. and the research leading to these results has received funding from the T. von Zastrow foundation, the FWF Wittgenstein award (Z 271-B19), the Austrian Academy of Sciences, the Innovative Medicines Initiative 2 Joint Undertaking (JU) under grant agreement No 101005026, and the Canada 150 Research Chairs Program F18-01336 as well as the Canadian Institutes of Health Research COVID-19 grants F20-02343 and F20-02015. Additionally, this project has received funding from the Innovative Medicines Initiative 2 Joint Undertaking (JU) under grant agreement no. 101005026. The JU receives support from the European Union’s Horizon 2020 research and innovation program and EFPIA.

## Author contributions

R.H.S. generated and developed the intervention concept, designed and analyzed experiments, and wrote the manuscript with input from all the authors. R.A.P. developed the intervention concept and selected the nebulizer for use in the experiments. S.K.L. developed the intervention concept. H.S.H. developed the intervention concept. N.R.W. generated and developed the intervention concept. P.T.W. generated and developed the intervention concept. P.S. planned and performed experiments and analyzed results. L.P. planned and performed experiments and analyzed results. R.G. planned and performed experiments and analyzed results. A.H. planned and performed experiments and analyzed results. S.K. planned and supervised experiments and analyzed results. D.B. planned experiments and analyzed results. J.M.W. planned and supervised experiments and analyzed results. Q.L. performed experiments and analyzed results. J.B. performed experiments and analyzed results. J.D.M. planned and supervised experiments and analyzed results. R.L.M. performed experiments and analyzed results. B.D.C. planned experiments and analyzed results. D.L.M. planned and supervised experiments and analyzed results. J.W.R. planned and conducted experiments and analyzed results. B.T.C. planned and conducted experiments and analyzed results. R.G. developed the intervention concept and analyzed experiments, S.H. developed the intervention concept and analyzed experiments, G.W. developed the intervention concept, designed and analyzed experiments, and provided clinical grade recombinant soluble human ACE2 (APN01). J.M.P. developed the intervention concept, helped with writing, and analyzed experiments.

## Competing interests

G.W., R.G. and S.H. are employed by Apeiron Biologics A.G. J.M.P was a founder of Apeiron, is a current shareholder and inventor of APN01. Other authors declare no competing interests.

## Supplemental Materials

**Supplemental Table 1.** ACE2 binding to SARS-CoV-2 Spike protein is not significantly altered after nebulization. All samples from trials 1-4 were normalized to 27 µg/ml according to R&D Systems ELISA measurements and assayed for binding to the SARS-CoV-2 Spike RBD. Data are recorded as EC50 values in ng/ml with 95% confidence intervals. The differences between Pre-Nebulization and Post-Nebulization binding activity are indicated as a % change for each APN01 dilution. In this Table, Pre-Nebulization corresponds to the Control samples in Figures 3 and 4 of the main text.

|  | Sample | EC50 | 95% CI |
| --- | --- | --- | --- |
| Trial 1<br>(2.5 mg/ml) | Pre-Nebulization | 58.41 | 40.56-85.50 |
|  | Unnebulized Volume | 41.08 | 34.85-48.60 |
|  | Post-Nebulization | 42.62 | 33.18-55.24 |
|  | % change Pre to Post | -27.03 |  |
| Trial 2<br>(2.5 mg/ml) | Pre-Nebulization | 44.62 | 39.42-50.58 |
|  | Unnebulized Volume | 43.65 | 37.55-50.85 |
|  | Post-Nebulization | 39.81 | 34.60-45.86 |
|  | % change Pre to Post | -10.78 |  |
| Trial 3<br>(0.1 mg/ml) | Pre-Nebulization | 57.32 | 43.41-76.55 |
|  | Unnebulized Volume | 50.05 | 44.29-56.70 |
|  | Post-Nebulization | 55.82 | 48.18-64.85 |
|  | % change Pre to Post | -2.62 |  |
| Trial 4<br>(0.1 mg/ml) | Pre-Nebulization | 43.81 | 38.53-49.89 |
|  | Unnebulized Volume | 34.94 | 32.28-37.79 |
|  | Post-Nebulization | 43.75 | 35.38-54.33 |
|  | % change Pre to Post | -0.14 |  |

**Supplemental Table 2.**
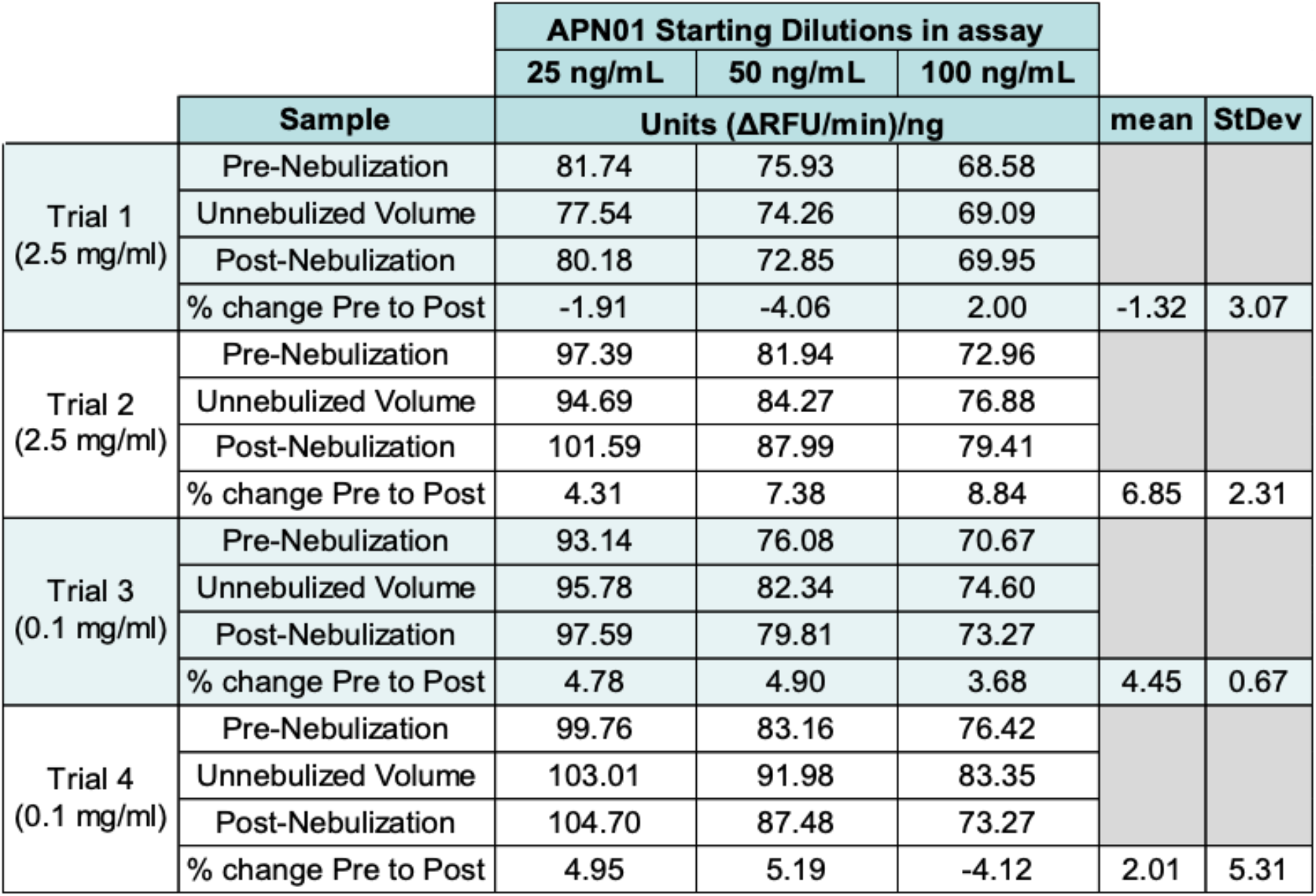
APN01 enzymatic function remains unaltered after nebulization. Data from trials 1-4 were recorded as (ΔRFU/min)/ng for the three starting dilutions of APN01 at 25 ng/ml, 50 ng/ml, 100 ng/ml. The difference between Pre-Nebulization and Post-Nebulization enzymatic activity is indicated as a % change for each APN01 dilution and the mean change and standard deviation across all three dilutions. In this Table, Pre-Nebulization corresponds to the Control samples in Figures 3 and 4 of the main text.

**Supplemental Table 3. Toxicology parameters studied\***.

A. Test Atmospheres
B. Inhaled Dose
C. Mortality and Clinical Signs
D. Body Weights and Body Weight Changes
E. Food Consumption
F. Clinical Pathology
G. Heart Rate and Blood Pressure
H. Ophthalmic Examination
I. Electrocardiography
J. Respiratory Function
K. Peripheral and Venous Blood Oxygen Saturation and Blood pH
L. Functional Observational Battery
M. Serum Drug Levels and Toxicokinetics
N. Organ Weights
O. Gross Pathology and Histopathology

*This study was conducted under Good Laboratory Practices. The table lists the parameters studied. Data for A, B and M are cited in the text. Detailed data (available on request) for the other parameters did not reveal treatment-related abnormalities.

**Supplemental Figure 1.**
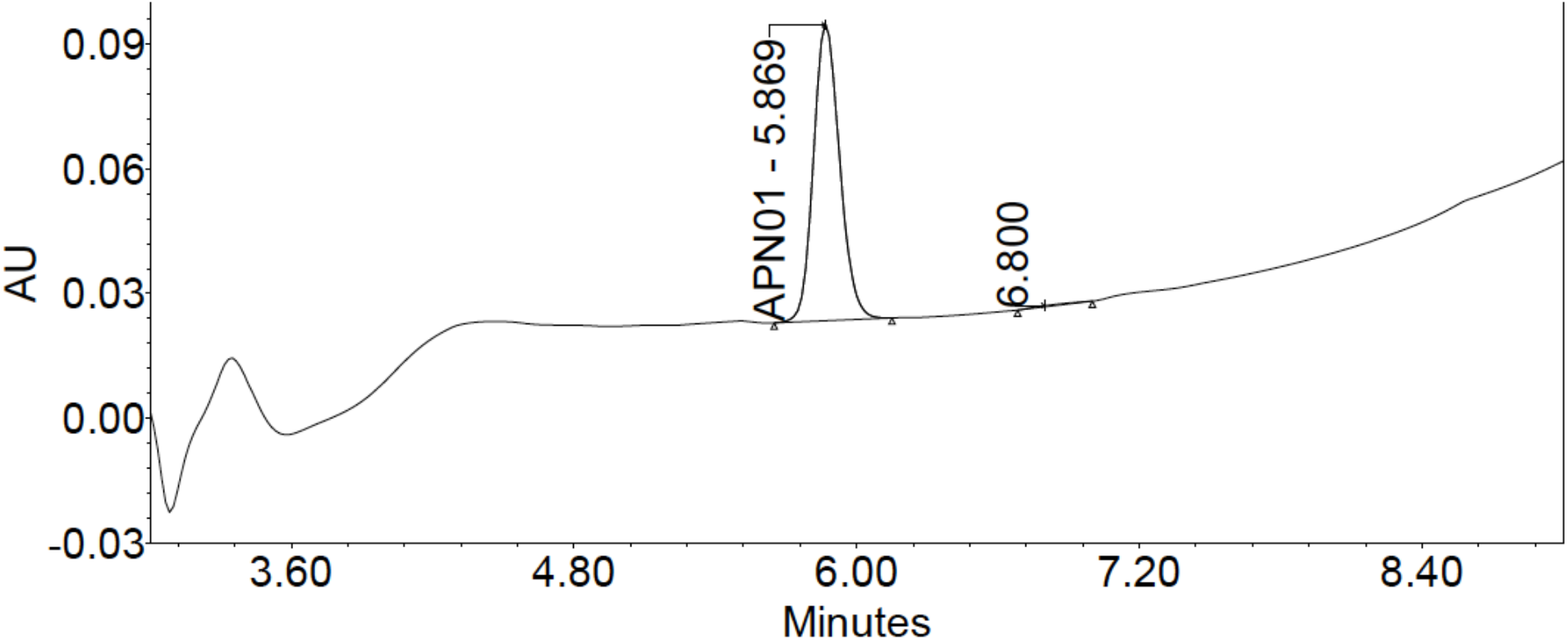
HPLC chromatogram illustrating quantitation of APN01 extracted from filters placed in the nebulized atmosphere at 0.075 mg/L APN01. Absorbance Units (AU) monitored at 220 nm are plotted as a function of time. Chromatography conditions are described in Materials and Methods. APN01 was resolved as a single peak eluting between five and six minutes. The concentrations of test article in the processed filter samples were determined from each sample’s peak area using the linear regression parameters derived from the calibration curves and correcting the resulting concentration by multiplying by the appropriate dilution factor, as applicable. In addition to supporting quantitation of APN01 in the nebulized atmosphere, this result supports maintenance of the physical integrity of APN01 during the process of aerosolization.

## Notes

### Competing Interest Statement

Gerald Wirnsberger, Romana Gugensberger, and Sonja Holler are, or were, employed by Apeiron Biologics A.G. Josef M. Penninger was a founder of Apeiron, is a current shareholder and inventor of APN01. Other authors declare no competing interests.

### Summary of Updates

This version corrects formatting problems introduced to the original submission on conversion to .pdf format.

